# Unipept in 2024: Expanding Metaproteomics Analysis with Support for Missed Cleavages, Semi-Tryptic and Non-Tryptic Peptides

**DOI:** 10.1101/2024.09.26.615136

**Authors:** Tibo Vande Moortele, Bram Devlaminck, Simon Van de Vyver, Tim Van Den Bossche, Lennart Martens, Peter Dawyndt, Bart Mesuere, Pieter Verschaffelt

## Abstract

Unipept, a pioneering software tool in metaproteomics, has significantly advanced the analysis of complex ecosystems by facilitating both taxonomic and functional insights from environmental samples. From the onset, Unipept’s capabilities focused on tryptic peptides, utilizing the predictability and consistency of trypsin digestion to efficiently construct a protein reference database. However, the evolving landscape of proteomics and emerging fields like immunopeptidomics necessitate a more versatile approach that extends beyond the analysis of tryptic peptides. In this article, we present a significant update to the underlying index structure of Unipept, which is now powered by a Sparse Suffix Array index. This advancement enables the analysis of semi-tryptic peptides, peptides with missed cleavages, and non-tryptic peptides such as those encountered in other research fields such as immunopeptidomics (e.g. MHC- and HLA-peptides). This new index benefits all tools in the Unipept ecosystem such as the web application, desktop tool, API and command line interface. A benchmark study highlights significantly improved performance in handling missed cleavages, preserving the same level of accuracy.

**For TOC Only:** 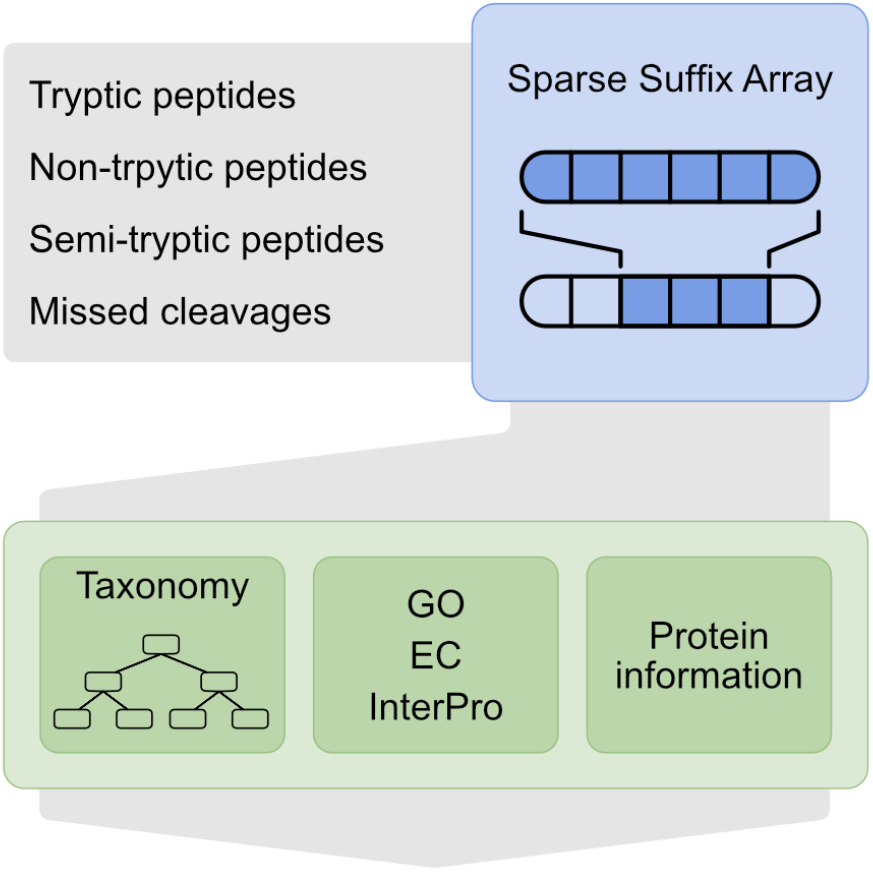

## Introduction

Unipept^1^ is an established ecosystem of user-friendly tools for the downstream analysis of metaproteomics data, initially launched as a web application in 2012 and expanded over the years to include a REST API^2^, a command line interface (CLI)^2^, and a desktop application^3^. All tools in the Unipept ecosystem depend on the same underlying API server, which can be either accessed via our public server or self-hosted for customized workflows.

Unipept excels in rapidly annotating a set of tryptic peptides with taxonomic and functional information. It accomplished this by utilizing a highly optimized relational database. For every tryptic peptide in UniProt^4^, Unipept’s database stored the precalculated taxonomic lowest common ancestor (LCA)^5^ and the linked functional annotations (EC^6^, GO^7,8^, InterPro^9^). For version 2024_03 of UniProt, this resulted in taxonomic and functional information on 1.3 billion tryptic peptides with a total database size of 1.875 TB. This required a server with at least 60 GB of memory to efficiently retrieve results. For example, it only took 11 seconds to analyze the SIHUMIx^10^ sample in the CAMPI benchmark dataset^10^ of 24424 tryptic peptides.

First and foremost, in metaproteomics, samples are commonly prepared by adding trypsin to cleave proteins at specific sites^11^. This is why constructing a precomputed index of all unique tryptic peptides from UniProtKB is feasible. From time-to-time, however, trypsin will miss a cleavage site, creating a so-called missed cleavage. By default, prior versions of Unipept assumed there are no missed cleavages present in a peptide, which means these cannot be matched to proteins. To solve this problem, a solution was retro-fitted to handle these missed cleavages. By enabling the “advanced missed cleavage” setting, Unipept scans the input peptide for missed cleavage positions, splits the peptides and performs separate lookups for each tryptic fragment. The resulting set of proteins are intersected, which results in a collection of possible protein matches, potentially containing the original peptide. Every protein in this collection is then scanned left-to-right to ensure that the original peptide is present, and not only its separate tryptic fragments. All these extra steps have a significant impact on performance. Enabling “advanced missed cleavage handling” increased the analysis time from 11 seconds to 6 minutes 33 seconds for processing the SIHUMIx dataset S07. In addition, the API and CLI did not support missed cleavage handling. Reviewing literature citing Unipept and our web server logs both reveal that almost all users enable missed cleavage handling, underscoring the importance of optimizing its performance. Although this feature significantly slows down the analysis, it has been widely adopted for its ability to enhance analysis accuracy.

Secondly, Unipept could not deal with semi-tryptic peptides (i.e. peptides where only one terminus is tryptic). This problem becomes particularly significant when proteolytic activity is less predictable, such as in the complex environment of the gut microbiome^12^.

Finally, support for matching arbitrary peptides such as non-tryptic peptides, which are often encountered in neighboring fields like immunopeptidomics^13^, was lacking in Unipept. Immunopeptidomics involves the analysis of HLA (human leukocyte antigen) peptides that play a crucial role in immune system functioning, as they are involved in presenting fragments of proteins (peptides) to T cells, enabling the immune system to distinguish between self and foreign antigens.

In this manuscript, we introduce a significant update to Unipept, consisting of a new index structure based on Sparse Suffix Arrays (SSA). This new index replaces the old relational database with precomputed data. Using this new index, Unipept can match arbitrary peptides (i.e. also semi-tryptic or non-tryptic peptides) instead of only tryptic peptides. In addition, it eliminates the performance penalty of missed cleavage handling. Because this index is used at the API-level of Unipept, these new features are automatically available to all tools in the Unipept ecosystem.

## Methods

In the Unipept ecosystem, components such as the Unipept Web application, Unipept Desktop app, CLI and third-party applications submit peptides to a central API for analysis. By making improvements to this API, all tools in the ecosystem automatically benefit. To support arbitrary peptide matching, the relational database in the Unipept API was replaced by a sparse suffix array (SSA), an index designed for efficient searching across multiple sequences. In addition, we completely rewrote the API itself from Ruby-on-Rails to Rust, a low-level, performance oriented programming language which achieves better performance and memory efficiency.

Peptides from a user sample are sent to the API, which in turn processes these peptides by querying the SSA index. This index matches them against all proteins in the UniProtKB database. Relevant taxonomic and functional annotations are retrieved from an optimized in-memory datastore, also implemented in Rust. The API then compiles this information and returns a list of proteins corresponding to each peptide, along with their associated taxonomic and functional annotations. These results are then passed back to the user (see Figure 1). Note that instead of relying on precalculated values, the taxonomic and functional annotations are now calculated on-the-fly.

**Figure 1:**
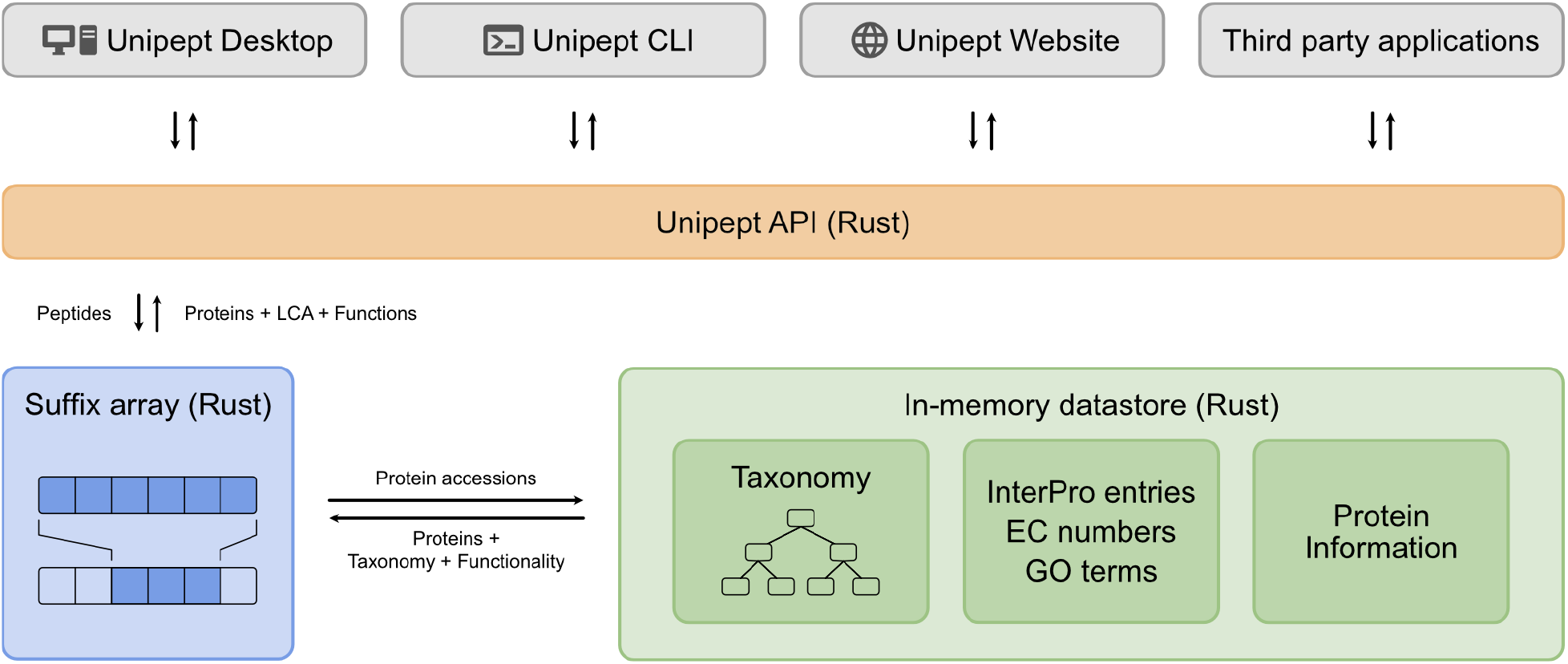
Current architecture of Unipept. Because each component of the Unipept ecosystem (in gray) relies on a common API (in orange) for their data analyses, improving this API immediately enables support for arbitrary peptide analysis and boosts performance for the entire ecosystem.

The SSA index organizes all suffixes of a given set of strings in sorted order, enabling fast retrieval of matching subsequences. Each suffix is stored as a 64 bit integer, pointing to its starting position in the text. Due to the sheer number of suffixes that must be referenced, the SSA index demands substantial memory. Therefore, we employed a number of techniques to reduce memory usage. These include sparseness (retaining only a subset of the suffix array without compromising accuracy), compression and bit packing. The following sections will explore these methods in detail, discussing their effects on both performance and memory efficiency.

### Constructing the suffix array

The SSA index is built by concatenating all protein sequences in the UniProtKB database. Given the vast number of proteins, this concatenated text and its corresponding suffix array can grow very large. Version 2024_04 of UniProtKB contains nearly 246 million proteins, totalling approximately 87 billion amino acids. Since each suffix is represented by a 64-bit integer, and the number of suffixes matches the number of amino acids, the resulting dense suffix array requires 64 × 87 × 10^9^ bits, equivalent to 696 GB of memory.

The suffix array itself can be constructed using the popular libdivsufsort^14^ or libsais^15^ libraries. Both implementations provide linear-time construction algorithms with a *5n + O(1)* memory complexity, where *n* is the size of the text being indexed. For version 2024_04 of UniProtKB, constructing the suffix array requires a machine with at least 750 GB of memory. Using the libdivsufsort library on a single core of a modern CPU (AMD EPYC 7773X), this process takes approximately 5 hours of computation time.

### Sparse suffix array

Although a dense suffix array provides optimal performance, it demands a substantial amount of memory. To decrease the amount of memory required for storing the suffix array, a sampling step can be introduced, resulting in a sparse suffix array (SSA). In an SSA, only every *k*-th suffix of the input text is retained, reducing the array’s memory footprint to 1/k^th^ of the original.

However, using a sparse suffix array introduces a limitation: an SSA with sparseness *k* only supports searching peptides that are at least *k* amino acids long, and overall lookup performance may decrease. Luckily, this compromise does not introduce a problem for Unipept, since it already restricted the search to peptides with a minimum length of 5 amino acids This aligns with limitations of mass spectrometery which struggles to reliably detect very short peptides. Moreover, such short peptides are prevalent in many proteins and tend to produce overly generic functional and taxonomic analysis results, offering limited information.

By using a sparseness factor of *k = 3*, we significantly reduce the memory requirements of the suffix array while preserving efficient search capabilities. From the original dense suffix array size of 696 GB, this approach reduces the memory usage to 696 / 3 = 232 GB, resulting in a much more manageable sparse suffix array.

### Suffix array compression

In addition to increasing the sparseness factor *k*, memory requirements can be further decreased by compressing the (sparse) suffix array. Each suffix represents a position in the text, and while the Rust programming language defaults to using 64 bits per suffix, this is unnecessarily large for the 87 billion amino acids in UniProt. To represent these 87 × 10^9^ different suffixes, we only need⌈log_2_(87 * 10^9^)⌉= 37 bits per suffix, eliminating 27 unused bits per suffix. By storing all suffixes sequentially in a custom bitarray, this compression significantly reduces memory usage.

For a sparse suffix array with *k = 3*, which initially requires 232 GB of memory, applying this compression reduces the size to just 133 GB—a 57% reduction in memory footprint. This optimization allows deploying the suffix array on servers with significantly less memory. Figure 2 illustrates the relative memory usage of each component within the final SSA index.

**Figure 2:**
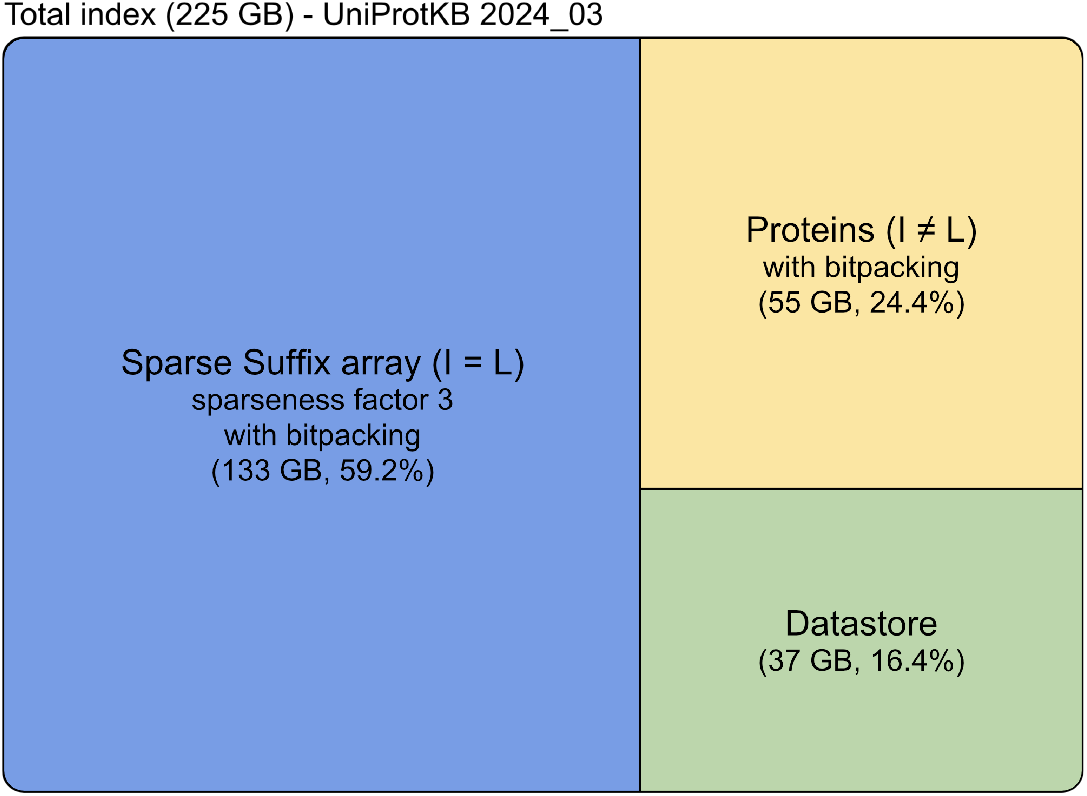
Memory footprint of the individual components in the SSA index built from UniProtKB version 2024_03.

### Equating Isoleucine and Leucine

In metaproteomics data analysis, the amino acids isoleucine (I) and leucine (L) are typically treated as equal. To support this requirement, the new index must natively accommodate the equivalence of I and L. Previous versions of Unipept achieved this by duplicating the entire database with I and L equated (I=L), effectively maintaining two versions of the index. However, applying this duplication strategy to the suffix array would double the memory requirements, making it impractical. Therefore, we only construct an index where every occurrence of leucine (L) is replaced with isoleucine (I).

When the “equate I=L” option is enabled, we similarly substitute leucine with isoleucine in the query peptides and directly return the results. For searches where isoleucine and leucine are treated as distinct, an additional filtering step is applied. In this step, each match from the initial search is verified by comparing the occurrences of I and L in the original, unmodified peptide with the corresponding amino acids in the unmodified protein sequences. Any mismatches where I and L were incorrectly equated are filtered out, ensuring only valid matches proceed.

### Maximal number of matches

A single peptide can potentially match thousands of proteins. Because we now aggregate all protein matches on demand, this can significantly slow down an analysis. Moreover, the results for such peptides are often quite generic because of their high occurrence. Out of all 1.3 billion tryptic peptides in UniProtKB only 0.001% result in more than 10 000 protein matches. For 95% of these extremely common peptides, the taxonomic lowest common ancestor^16^ corresponds to the root node of the NCBI taxonomy (as detailed in the supplementary data S1). Rather than explicitly computing the lowest common ancestor, we opted to heuristically assign the root taxon to all peptides with more than 10 000 matches. This heuristic greatly improves performance while causing only a minimal reduction in accuracy - in UniProt 2023.3, only 639 out of 1.3 billion tryptic peptides are assigned a more generic LCA under this method.

## Results

Replacing the relational database with our novel SSA index should have minimal impact on the accuracy of the analysis results. However, we anticipate a significant performance improvement, particularly when missed-cleavage handling is enabled. To evaluate this impact, we analyzed samples from the highly used SIHUMIx benchmark dataset^10^ using the Unipept web application. Version 5.0 of Unipept relies on the old relational database, whereas version 6.0 employs the new SSA index.

### Accuracy comparison

To assess the impact on accuracy, we used sample S07 from the SIHUMIx dataset containing 24424 peptides. The full results can be found in supplementary data S2. In the analysis with Unipept 5.0, 297 peptides (1%) could not be matched. In contrast, Unipept 6.0 successfully matched every peptide in the sample. When examining the taxonomic annotations for each peptide, we observed that in Unipept 5.0 2,680 peptides were annotated with an LCA at the species level, whereas this number slightly decreased to 2,551 for Unipept 6.0.

The slight shift in results arises from a fundamental difference in how the relational database and the SSA index handle peptide queries. In the relational database, peptides were assumed to be tryptic, implicitly incorporating information about the amino acids preceding and following the peptide. In contrast, the SSA index can match arbitrary peptides, so this implicit context is no longer considered during matching, which may result in additional protein matches. Given the minimal impact of these differences, we have decided not to implement some kind of tryptic emulation of the old database.

### Performance comparison

To evaluate performance, we analyzed six tryptic peptide samples from the SIHUMIx data set. Each sample contained around 25 000 peptides, including both tryptic peptides and peptides with missed cleavages. Duplicate peptides were not filtered and the “equalize IL” option was enabled. Figure 3 shows wall clock times for searching these samples against Unipept 5.0 with and without missed cleavage handling, as well as Unipept 6.0, which natively supports missed cleavage handling.

**Figure 3:**
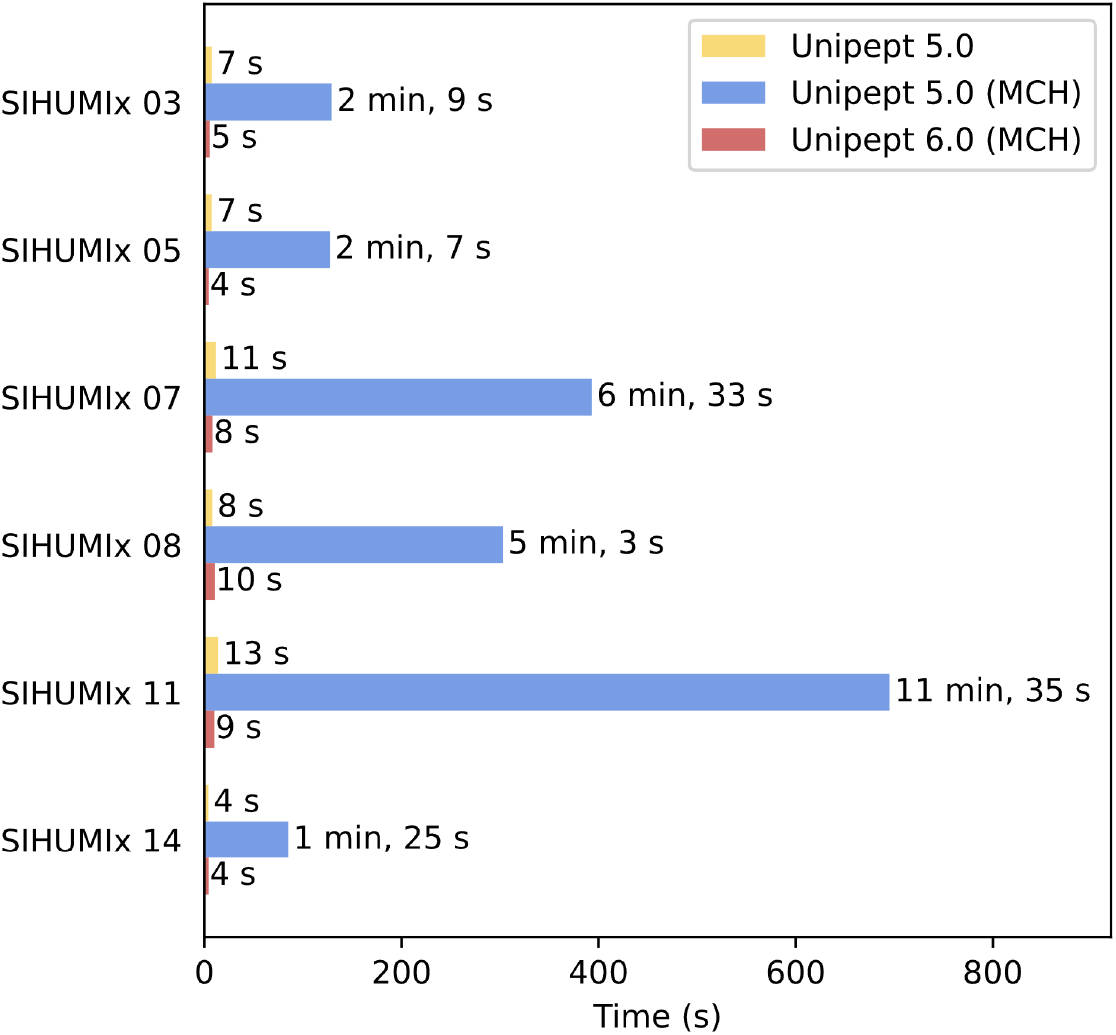
Execution times for Unipept 5.0, and Unipept 5.0 with missed cleavage handling (MCH) enabled and Unipept 6.0. Each SIHUMIx sample contains approximately 25000 tryptic peptides. All analyses used the I = L option.

As a general observation, Unipept 5.0 without missed cleavage handling processes peptides at the same speed as Unipept 6.0 with missed cleavage handling. This represents a 20-70 times speed improvement compared to using missed cleavage handling using Unipept 5.0. For example, when analyzing CAMPI^17^ dataset S07, the search time with missed cleavage handling dropped from 6 minutes 33 seconds using Unipept 5.0 to just 8 seconds with Unipept 6.0. This is comparable to the 11 seconds it took to analyze the dataset using Unipept 5.0 without missed cleavage handling.

### Comparison with other tools

Tools such as the UniProt peptide search tool^18,19^, the Expasy ScanProsite tool^20^ and Unipept all provide functionality for matching peptides against the UniProtKB database. In this section, we compare the feature sets and performance of these tools with Unipept 6.0. Because of the poor performance of some of these tools, our tests were restricted to a single, randomly selected tryptic peptide: ISPAVLFVIVILAVLFFISGLLHLLVR.

The features and execution times of all tools are available in Table 1. Both the UniProt peptide search tool and Unipept 6.0 provide similar features, including the ability to use the I=L option and perform searches across the entire UniProtKB database. However, their performance differs significantly: while the UniProt peptide search tool completes searches in a few seconds to minutes, Unipept 6.0 completes the same task in under a few milliseconds. The Expasy ScanProsite tool adopts a different approach, offering a broad array of approximate matching options. However, its search is limited to a subset of UniProtKB, specifically proteins from reference genomes, which account for only about one-third of the entire UniProtKB database. Searching for the reference peptide with ScanProsite took 5 minutes. Unipept 5.0 and Unipept 6.0 both support searching the entire UniProtKB database and allow the I=L option, performing equivalently in terms of accuracy. However, Unipept 5.0 is restricted to tryptic peptides, whereas Unipept 6.0 does not have this limitation.

**Table 1:** Comparison of Unipept 5.0, Unipept 6.0, UniProt peptide search (UPS) tool and ExpasyScanProsite (EPS) tool. [IL] in the approximate matching row indicates only I and L can be equated, while ∼ tryptic in the searchable peptides row indicates only tryptic peptides or tryptic peptides with missed cleavages can be found.

|  | UPS | ESP | Unipept 5.0 | Unipept 6.0 |
| --- | --- | --- | --- | --- |
| Used proteins | all | Ref. proteomes | all | all |
| Approximate match | [IL] | flexible | [IL] | [IL] |
| Execution time | 1 s - 20 min | 5 min | **< 5 ms** | **< 5 ms** |
| Searchable peptides | all | all | ~ tryptic | all |

## Conclusions

In this manuscript, we present a significant update to the underlying index structure of Unipept. This index is implemented in Rust and allows fast on-demand calculation of both taxonomic and functional annotations for a set of peptides. Because data processing no longer relies on precalculated results for tryptic peptides, Unipept can now match arbitrary peptides. This allows us to perform “missed cleavage handling” by default up to 70 times faster, making it feasible to integrate missed cleavage handling into the API, CLI, and third-party tools—a capability that was previously unavailable.

Looking ahead, Unipept is well-positioned to continue evolving, with a robust foundation for migrating features currently exclusive to the desktop application onto the web platform. One prominent example is the potential introduction of an ad-hoc filtering feature for UniProtKB, similar to the custom database construction already available in the Unipept Desktop app. This functionality would allow researchers to query a targeted subset of proteins within the extensive UniProtKB database, leveraging taxonomic data such as results from prior metagenomic experiments. By transitioning these advanced features to the web, Unipept can enhance accessibility for a wider range of users, eliminate the need for local software installations, and improve the overall user experience.

## Acknowledgements

PV acknowledges funding by Ghent University [BOF/01P10623]. L.M. acknowledges funding from the Research Foundation Flanders (FWO) [G028821N] and [G010023N]. TVDB acknowledges funding by the Research Foundation Flanders (FWO) [1286824N].

## Data Availability Statement

Unipept is available as open-source software, with its source code accessible on GitHub. The SIHUMI samples analyzed in this study are publicly available through the CAMPI dataset. Additional data files related to this study are provided as supplementary data.

Supporting information

The following supporting information is available free of charge at ACS website:

- Supplementary Data S1. In-depth analysis of tryptic peptides and associated UniProtKB proteins
- Supplementary Data S2. Data validation files

